# Highly Efficient Lentiviral Transduction of Human iPSC-Derived Microglia and Macrophages

**DOI:** 10.64898/2026.05.23.727402

**Authors:** Srilakshmi C. Goberdhan, Magdalena A. Czubala, Sophie E. Thomas, Philip R. Taylor, Natalie Connor-Robson

## Abstract

**Background:** Microglia have become a cell type of interest in the neurodegenerative field given both genetic and pathological evidence for their role in disease development and progression. There has been a rapid growth of studies using iPSC-derived microglial models to understand the molecular mechanisms driving these neurological diseases. However, it remains difficult to transduce myeloid cells effectively which is critical when aiming to study the role of disease associated genes and pathways. Current methods require exposure to multiple viruses which is not suitable for all experimental paradigms. We have therefore sought and characterised a high efficiency promoter and plasmid design to allow high transduction efficacy with a single lentivirus.

**Results:** Using the spleen focus-forming virus (SFFV) promoter in combination with central polypurine tract (cPPT) and Woodchuck hepatitis virus post-transcriptional regulatory element (WPRE) plasmid elements gave significantly higher transduction efficiency and transgene expression than was achieved with commonly used promoters CMV and EF1α. This could then be further improved if required to over 90% transduction efficiency with the removal of lentivirus restriction factor SAM and HD domain-containing protein 1 (SAMHD1) by adding VPX.

**Conclusions:** Our findings allow for a simpler, more efficient and streamlined approach to transgene expression in iPSC-derived microglia and macrophages using only a single lentivirus. This minimises potential unintended side effects such as additional cellular activation and increased cell death.

## Background

Microglia are the main resident immune cells of the central nervous system (CNS), critical in brain homeostasis, immune defence and regulating neuroinflammation (1). The multifaceted role of microglia has been further elucidated through the identification of microglial states based on their transcriptional profiles including ‘homeostatic’, ‘lipid processing’, ‘antiviral’ and various inflammatory signatures (2). There is growing evidence of microglial dysfunction across the neurodegenerative spectrum including in Alzheimer’s Disease (AD) where microglia become reactive and cluster around the pathological hallmark of AD, amyloid beta plaques (3). Microglia have also been demonstrated to adopt a protective pro-inflammatory state which, when sustained, can result in neuron and/or myelin degeneration observed in Parkinson’s disease (4), amyotrophic lateral sclerosis (ALS) (5) and multiple sclerosis (MS) (6). Additionally, growing genetic evidence from genome-wide association studies (GWAS) demonstrates microglial disease associated genes in various neurodegenerative diseases (7–9). Comprehensive examination of microglial function and their disease related biology requires efficient manipulation of these genes of interest. These gene-targeted approaches will enable greater understanding of these critical cells, particularly when implemented in a high-throughput, human disease relevant model system like human induced pluripotent stem cell (iPSC) derived microglia.

Microglia and macrophages are challenging cell types to transduce with either adeno-associated virus (AAV) or lentiviral transduction methods and, even when successfully transduced, transgene expression is often low (10–14). Lentiviral transduction is advantageous given the larger transgene carrying capacity and stable expression however, human stem cell models, particularly microglia, have proved difficult to gain efficient lentiviral transduction, partly due to gene silencing over time (15). Previous work has shown the choice of promoter to be important to overcome this and should be carefully selected for the cell type being studied (16–18). Due to their innate function as immune cells, microglia express toll-like receptors (TLRs) and other anti-viral mechanisms, such as restriction factors (19) which allow them to sense and protect the CNS from viral infection. Therefore, typical lentiviral transduction has been difficult to apply in these cells and results in very low transduction efficiency which hinders experimental utility. Previous work has used co-transduction with additional factors including VPX viral-like particles to target these restriction factors to improve lentiviral transduction of microglia, but this requires transduction with multiple viruses and is not always effective (20).

In this study, we demonstrate that implementing the spleen focus-forming virus (SFFV) promoter alongside central polypurine tract (cPPT) and Woodchuck hepatitis virus post-transcriptional regulatory element (WPRE) into the plasmid backbone is sufficient to achieve highly efficient transduction with a single virus in iPSC-derived microglial and macrophage models which, if required, can be further enhanced with VPX. This allows highly efficient and stable expression of genes of interest across myeloid cell types, as well as offering a simplified workstream through the need for only a single virus in comparison to current methods. These improvements allow easier manipulation of gene expression in iPSC-derived microglia and macrophage models providing a useful system to study the molecular mechanisms of health and disease *in vitro*.

## Materials and Methods

### hiPSC culture and maintenance

The KOLF2.1 iPSC cell line was used throughout this work as it has been widely adopted as a reference line in the neurodegenerative field (21). iPSCs were cultured on hESC-Qualified Matrigel (Corning) with mTeSR™ Plus (Stem Cell Technologies 100-0276) and split as required with 0.5 mM ethylenediaminetetraacetic acid (EDTA) (Gibco 15575020) (22).

### Microglial/Macrophage Differentiation and cell characterisation

Microglia were differentiated and maintained in monoculture as previously described (23). Briefly, iPSCs were plated into an AggreWell™800 (Stem Cell Technologies) with EB media (mTeSR Plus base media with 50ng/mL BMP4 (PeproTech), 50ng/mL VEGF (PeproTech) and 20ng/mL SCF (PeproTech)) to form embryoid bodies (EBs). After 2 days, EBs were transferred to CELLSTAR® cell-repellent 6-well plates with an excess of media. After 6 days, EBs were transferred to T175 flasks with Factory Media (X-Vivo base media with 100 ng/mL M-CSF (PeproTech), 25 ng/mL IL-3 (PeproTech). After 4 weeks, the flasks were harvested for macrophage precursors which were differentiated into microglia (Advanced DMEM/F12 (Gibco) base media with N2 supplement (Thermo Fisher), 100 ng/mL IL-34 (PeproTech) and 10 ng/mL GM-CSF (Gibco)) or macrophages (X-Vivo base media with 100ng/mL M-CSF) according to the final plating media. In all experiments, microglia or macrophage precursors were terminally differentiated for a total of 10 days, a typical timepoint for experiments with these cell types and differentiation protocols.

### Immunocytochemistry and live imaging

Both microglia and macrophages were fixed with 4% paraformaldehyde and blocked using 10% normal donkey serum (Abcam ab7475-25ml) in 0.04% PBS-Tween (PBS-T), followed by the addition of primary antibodies (diluted in blocking solution) overnight at 4°C. Secondaries were added at 1:1000 in PBS-T for 2 hours prior to imaging on the Opera Phenix high throughput confocal microscope (Revvity). Both cell types expressed their respective markers – IBA1 (Abcam ab5076 *1:250*) and PU.1 (Cell Signalling Technology 89136S *1:100*) for microglia and CD45 (Proteintech 65109-1-IG-100UG *1:100*) and CD163 (Proteintech 16646-1-AP-20UL *1:100*) in macrophages as shown in in Figure 1.

**Figure 1:**
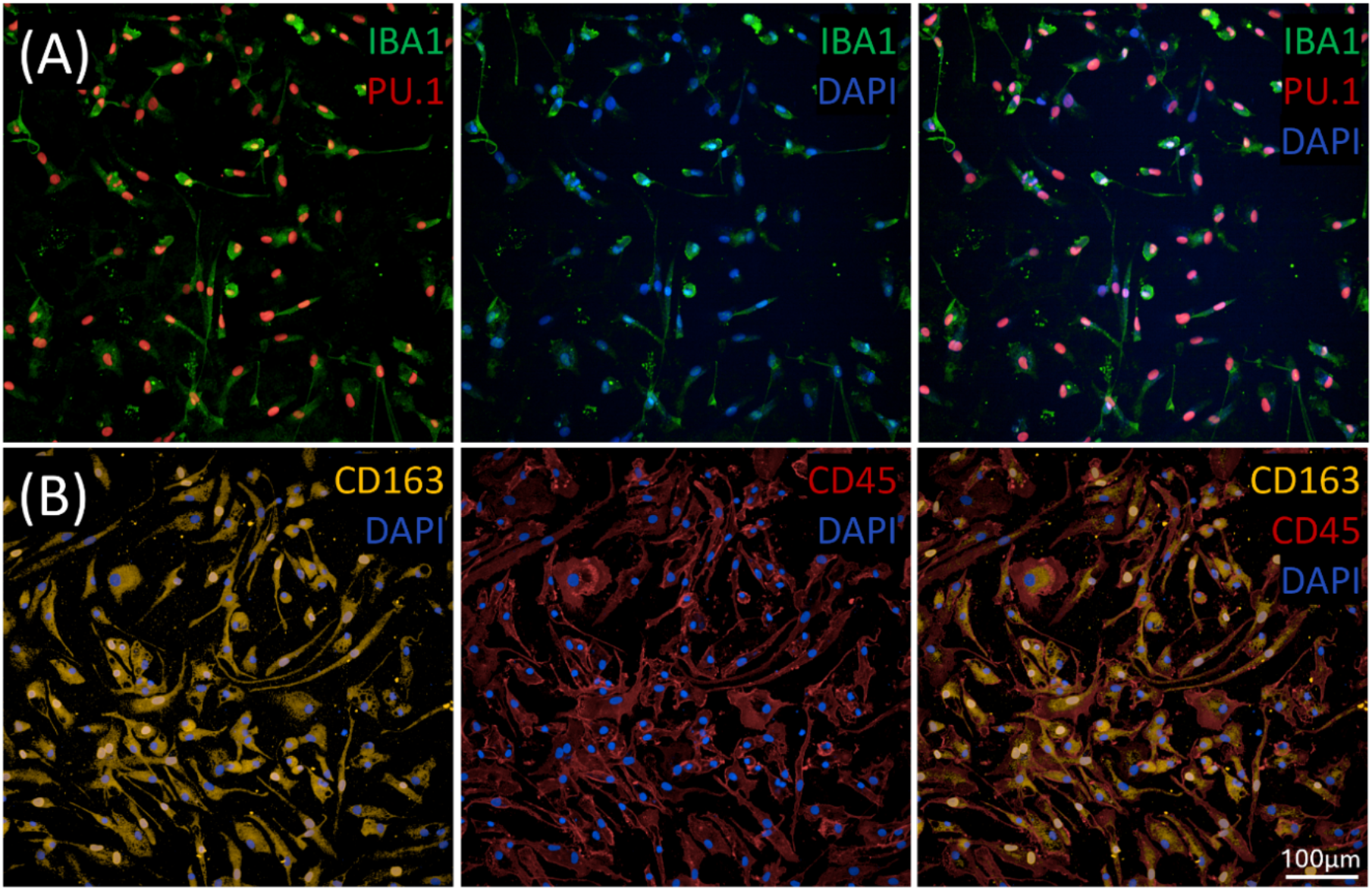
Confirmation of microglia and macrophage identity. **(A)**Representative images of immunostaining for microglia with IBA1, PU.1 and DAPI. **(B)**Representative images of immunostaining for macrophages with CD163, CD45 and DAPI. Scale bar represents 100µm.

Phagocytic activity was measured by the uptake of pHrodo Zymosan (Invitrogen P35364) through live imaging. In brief, all cells were stained with NucBlue LiveReady Probes (Molecular Probes R37605) and CellTracker Deep Red (Invitrogen C34565), and negative control wells treated with Cytochalasin D (BioTechne 1233/1) for 45 minutes prior to adding pHrodo red Zymosan at 50µg/mL and imaged every 20 minutes for up to 3 hours using the Opera Phenix.

### Western blot

To assess SAMHD1 suppression by VPX, microglia were plated at 1×10^6^ per well of a 6-well plate (Greiner) and treated with VPX at various time intervals. Cells were lysed with cold RIPA buffer (50 mM Tris at pH8.0, 150 mM NaCl, 1% NP-40/IGEPAL, 0.5% NaDeoxycholate, 0.1% SDS) with added cOmplete™, EDTA-free Protease Inhibitor Cocktail (Roche). Protein concentration was determined using the Pierce™ BCA Protein Assay Kit (Thermo Scientific) according to manufacturer’s instructions. 20 µg of each sample was loaded into a Novex WedgeWell™ 8 to 16% Tris-Glycine Mini protein gel run with 1X Novex Tris Glycine SDS Running Buffer and then transferred onto a PVDF membrane using the iBlot 2 system. Membranes were blocked in 4% milk prepared in 1X Tris-buffered saline (TBST, NaCl and Tris HCl in MilliQ + 0.1% Tween20) before SAMHD1 (12586-1-AP Proteintech) and GAPDH (ab8245 Abcam) primary antibodies were applied at 1:1000 each in 4% milk and incubated at 4°C on a roller overnight. Membranes were washed with TBST and incubated in secondary antibodies at 1:5000 in 4% milk, Goat anti-Mouse DyLight™ 800 (Invitrogen SA5-10176) and Donkey anti-Rabbit Alexa Fluor™ Plus 680 (Invitrogen A32802) for 1 hour at room temperature, protected from light. After TBST washes, the blot was imaged on the LI-COR Odyssey® DLx and quantified using the Image Studio Lite 5.2.5 software.

### Vector details and titration

All plasmid details are listed in Table 1. Plasmid A was received as a packaged and concentrated lentivirus from VectorBuilder (#VB010000-9492agg (24)). VPX (Plasmid D) was packaged as described previously (20) with modifications. In brief, HEK293T cells were transfected using the CalPhos Mammalian Transfection Kit (Takara 631312) with the packaging plasmid, pMD.2G (Table 1B) which encodes the VSV-G envelope protein. Prior to collection, media was changed from complete media (DMEM, high glucose, GlutaMAX Supplement, pyruvate (Gibco 10569010) base media with 10% Fetal Bovine Serum (FBS) (Gibco A5256701)) to Opti-MEM Reduced Serum Media (Gibco 31985070).

**Table 1.**
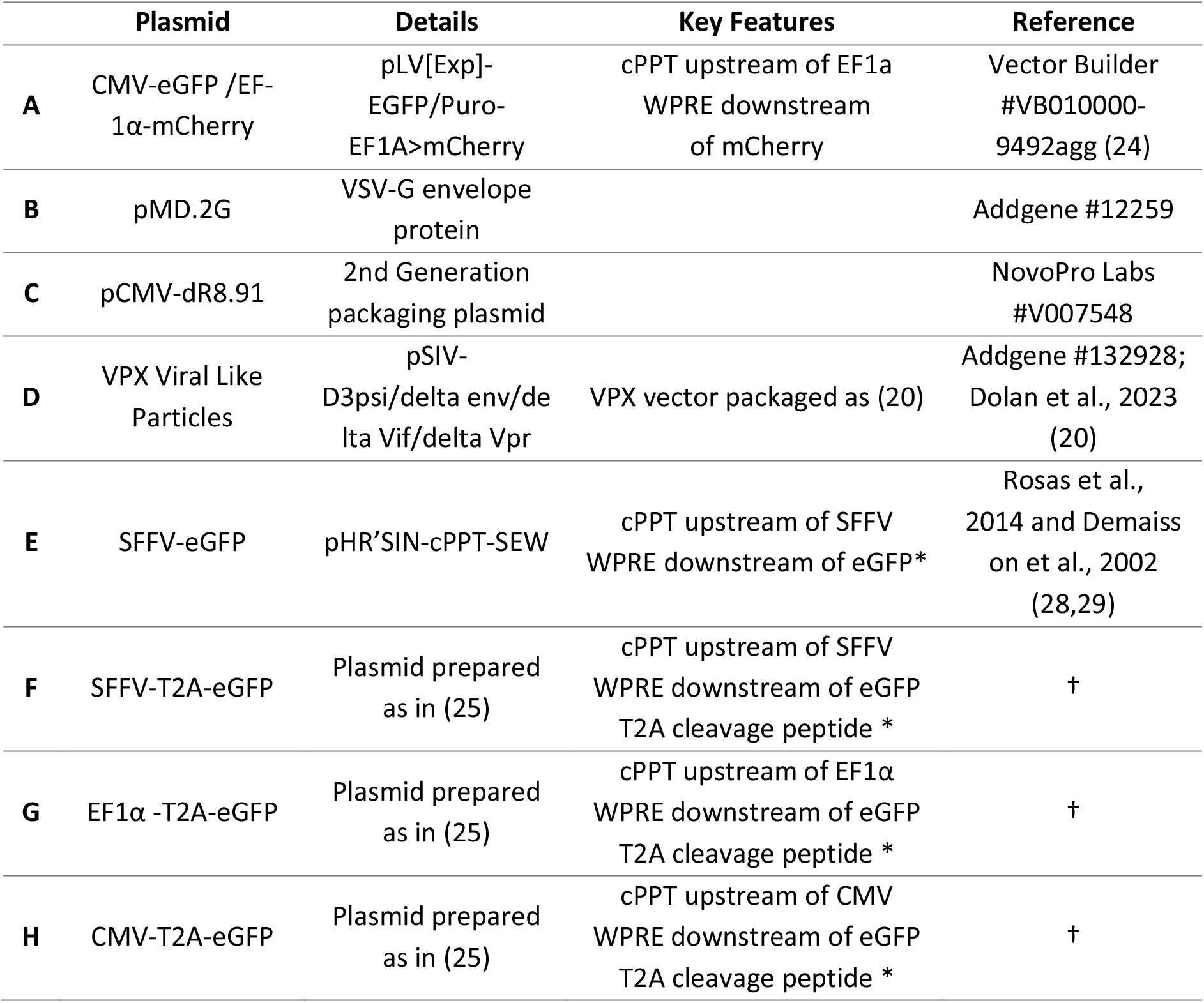
Details about plasmid vectors used in this study. *Vector maps in Supplemental Figure 1, † Plasmids available on request.

The SFFV promoter vector was pHR’SIN-cPPT-SEW plasmid (Table 1E) as previously described (25). Plasmids F-H have identical plasmid backbones, whereby the only difference is the promoter (Supplemental Figure 1). Sequences of EF1α (Plasmid G) and CMV (Plasmid H) promoter regions were cloned exactly as in Plasmid A. All three plasmids were packaged the same (25,26). Briefly, lentivirus was produced from each of the three plasmids using second generation packaging plasmids pMD.2G (Plasmid B) and pCMV-R8.91 (Plasmid C) and the Effectene transfection reagent (Qiagen 301425). Notably, all four lentiviruses (E-H) were concentrated using a 20% sucrose solution with ultracentrifugation to avoid FBS concentration as the viruses were used on cell types likely to become activated by components in FBS. Lentiviral titres were determined by quantitative PCR (qPCR) as previously described (27) to determine the multiplicity of infection (MOI).

### Microglia transduction

All lentiviral transductions were performed at MOI 30. Upon harvesting from the T175 flasks on DIV 0, cells were transduced with the lentivirus with or without VPX as illustrated in Supplemental Figure 2. VPX was used at 1μL/1×10^3^ cells as previously described (20). For the 24-hour VPX pre-treatment, cells were treated with VPX on DIV 0 and then lentivirus was added 24 hours later. In all three conditions, cells were left for 48 hours with lentivirus to ensure ample time for successful lentiviral infection prior to full media change to fresh microglial or macrophage media and matured until DIV10.

On DIV10, the nuclei of the transduced microglia/macrophages were live stained with NucBlue LiveReady Probes as per manufacturer’s instructions and imaged on the Opera Phenix high content confocal microscope on 20x water objective and image analysis was performed using Signals Imaging Artist 1.4.3 (Revvity). A stringent intensity threshold was implemented during image analysis to ensure only transduced cells were selected for analysis. Transduction efficiency was measured as the number of eGFP or mCherry-positive cells as a percentage over all the DAPI-positive cells. Mean fluorescence intensity (MFI) was defined as the intensity of eGFP/mCherry signal of the identified eGFP/mCherry positive cells as a mean per well. All experiments presented were repeated over three independent experiments with 3-4 technical replicates per experiment.

### Statistics

GraphPad Prism v11.0.0 (GraphPad) was used to generate graphs for statistical analysis. P values indicated in figure legends, * for *p* < 0.05, ** for *p* < 0.005, *** for *p<0*.*0005*, **** for *p* < 0.0001. Two-way ANOVA was used to investigate the effect of promoter and VPX treatment, except for Fig. 3B in which a one-way ANOVA was used to examine SAMHD1 expression over time.

**Figure 2:**
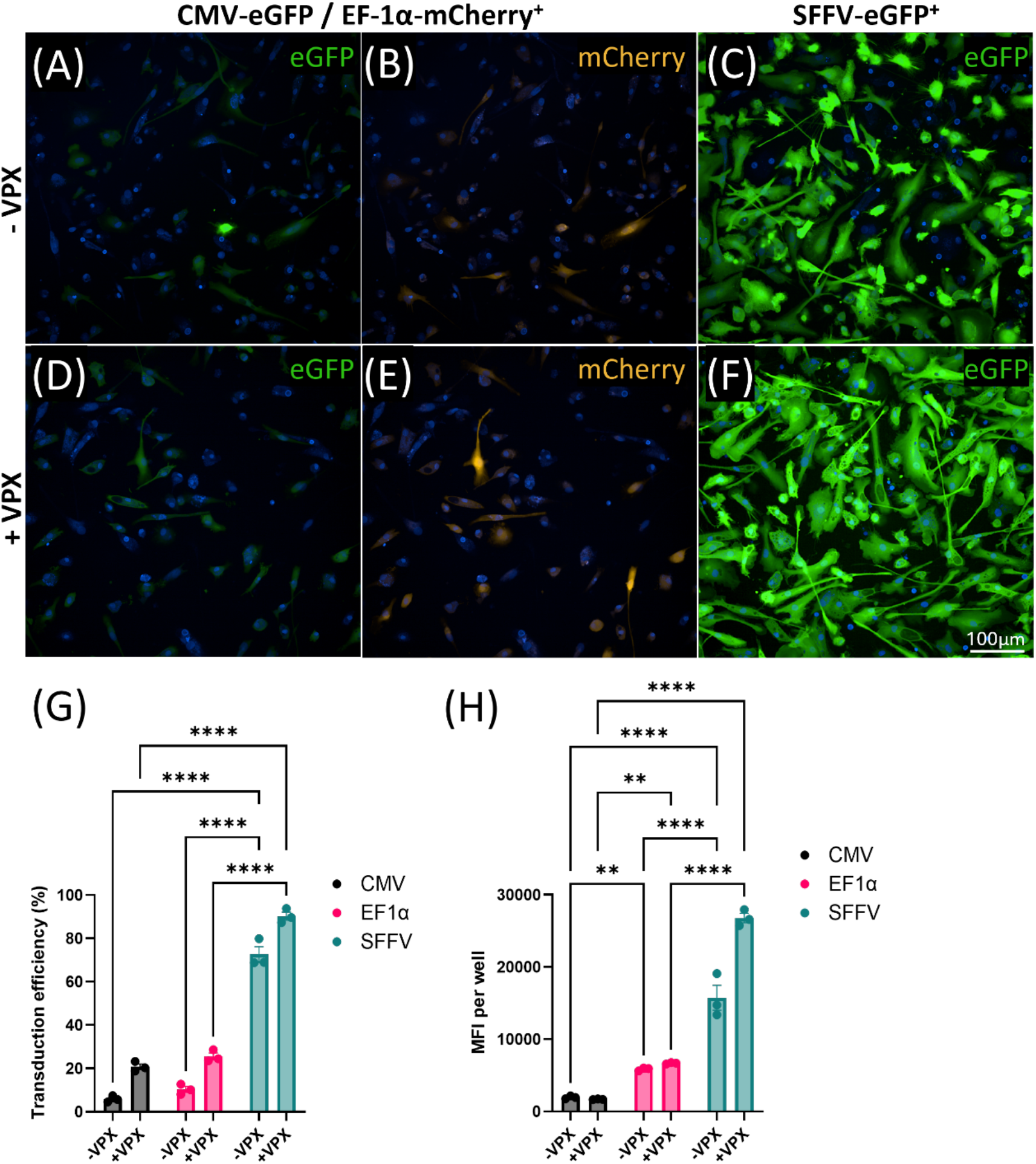
Effect of promoter on transduction efficiency in iPSC-derived microglia. Representative images of KOLF2.1 iPSC-derived microglia transduced with Plasmid A **(A)** CMV-eGFP, **(B)** EF1α-mCherry or Plasmid B **(C)** SFFV-eGFP in the absence of VPX or with VPX and Plasmid A **(D)** CMV-eGFP, **(E)** EF1α-mCherry or Plasmid B **(F)** SFFV-eGFP; scale bars represent 100µm. Quantification of **(G)** transduction efficiency: Two-way ANOVA (effect of promoter and VPX p=<0.0001), with Tukey’s post hoc test and **(H)** mean fluorescence intensity (MFI): Two-way ANOVA (effect of promoter and VPX p=<0.0001), with Tukey’s post hoc test. **p<0.005; ****p<0.0001; n=3 wells per condition; error bars denote mean ±SEM.

**Figure 3:**
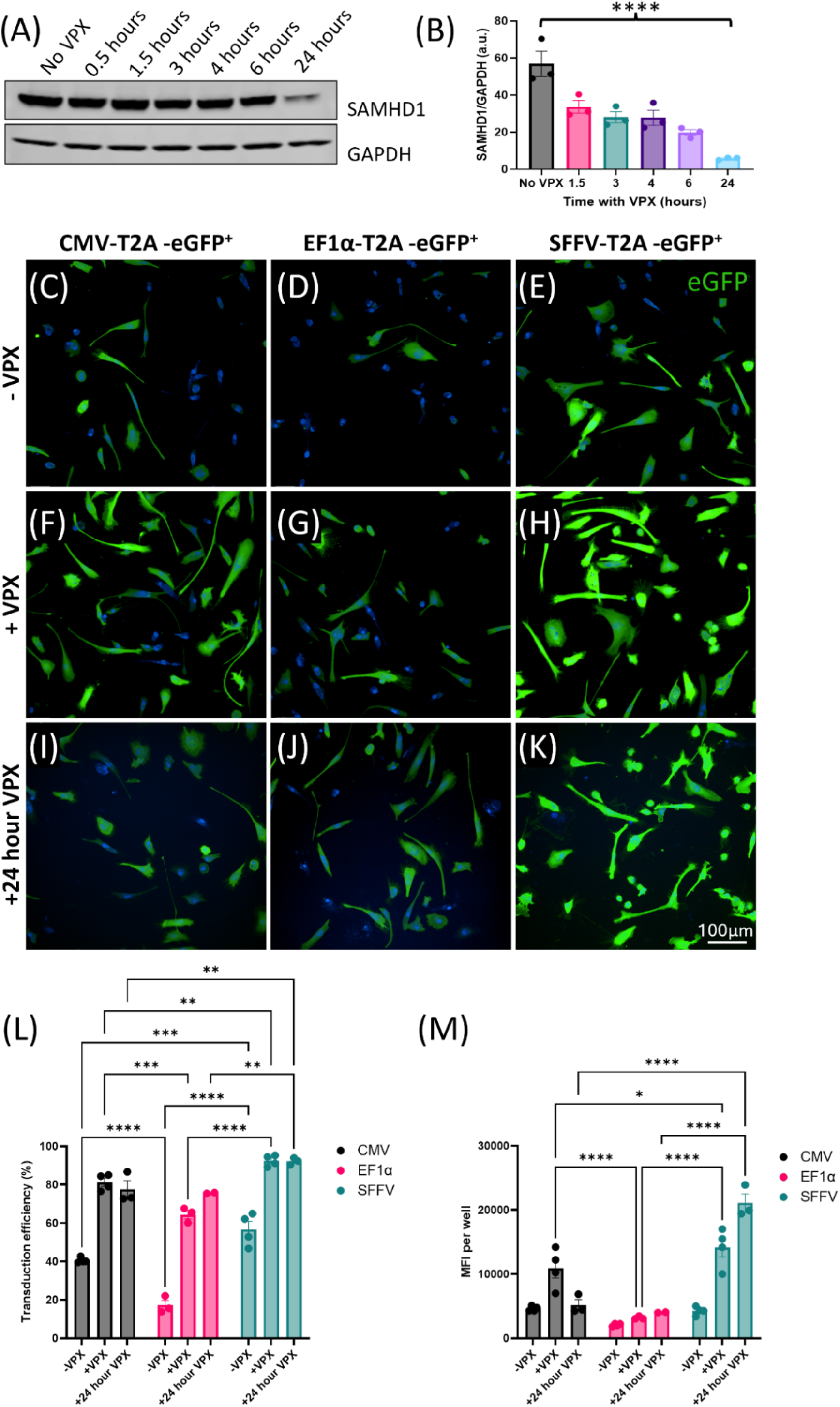
Comparing promoters in the same plasmid backbone confirms SFFV improves transduction efficiency which can be further enhanced with VPX. **(A)** Microglia were treated with VPX for 24 hours and harvested at various intervals, SAMHD1 depletion was measured by Western blot with **(B)** quantification of SAMHD1 protein normalised to GAPDH; one-way ANOVA ****p <0.0001. Representative images of iPSC-derived microglia transduced with **(C-E)** a single lentivirus, **(F-H)** the same lentivirus combined with VPX or **(I-K)** the same lentivirus of interest after a 24-hour pretreatment with VPX; scale bar represents 100µm. Quantification of **(L)** transduction efficiency: Two-way ANOVA (effect of promoter and VPX p=<0.0001), with Tukey’s post hoc test and **(M)** mean fluorescence intensity: Two-way ANOVA (effect of promoter and VPX p=<0.0001), with Tukey’s post hoc test. *p<0.05, **p<0.005, ***p <0.0005 and **** p<0.0001. n = 2-4 technical replicates per condition; error bars denote mean ±SEM.

## Results

### Using the SFFV promoter increases transduction efficiency

Previous work in the iPSC field has demonstrated higher transduction efficiency and transgene expression stability with the EF1α promoter compared to CMV (17). To determine if choice of promoter could improve transduction efficiency of human iPSC-derived microglia we compared CMV, as one of the most common ubiquitous promoters in mammalian cells for driving expression, EF1α as a constitutive promoter of human origin that has previously been reported as the most stable promoter in iPSC models (17) and finally, the SFFV promoter given its previous reported use in myeloid cells (25,30).

KOLF2.1 iPSC-derived microglial precursors were seeded at 1×10^3^ cells/mm^2^ and transduced at MOI 30 with either the VectorBuilder lentivirus gene expression control virus (24) in which the EF1α promoter drives mCherry expression and the CMV promoter drives eGFP (Plasmid A), or an in-house produced lentivirus in which the SFFV promoter drives eGFP expression (Plasmid E). The CMV and EF1α promoters were only able to drive detectable expression in 6% and 10% of the microglia, respectively, whereas the SFFV promoter was able to attain a transduction efficiency of 73% (Figure 2).

Transduction of microglia simultaneously with VPX and the plasmids outlined above demonstrated improvement in transduction efficiency as has been previously reported (20). This increased transduction efficiency to 21% for CMV, 26% for EF1α and 90% for SFFV. However, this increase was less substantial than that achieved through the change of promoter. We also assessed the mean fluorescence intensity for all conditions of the transduced cells (Figure 2H). The SFFV plasmid drove the highest mean fluorescence intensity (MFI) when the different promoters were used alone and the addition of VPX further increased MFI which was not evident with the CMV or EF1α plasmids.

### Examining the effect of the SFFV promoter in isolation in microglia

Although the results above demonstrate the lentiviral vector with the SFFV promoter is seemingly advantageous for driving expression in microglia, the use of different plasmid backbones (Table 1) likely meant that other vector elements could influence the transduction efficiency. In particular, the central polypurine tract (cPPT) and Woodchuck hepatitis virus post-transcriptional regulatory element (WPRE) have previously been shown to work in tandem to improve transduction efficiency by initiating DNA synthesis and enhancing gene expression post-transcriptionally (31– 36). To understand if these plasmid elements were beneficial, the sequences of the CMV and EF1α promoters used above (plasmid A) were cloned into the SFFV-T2A-eGFP (plasmid F), which uses the same backbone as SFFV-eGFP (plasmid E), replacing the SFFV promoter to create Plasmids G and H. In doing this we were able to standardise the plasmids as plasmid A had multiple promoters, ORF-stuffers and IRES sequences which may reduce transduction efficiency. These standardised plasmids were each packaged and concentrated concurrently as previously described (26). KOLF2.1 iPSC-derived microglia (Figure 3) and macrophages (Figure 4) were then transduced at MOI30 with and without VPX.

**Figure 4:**
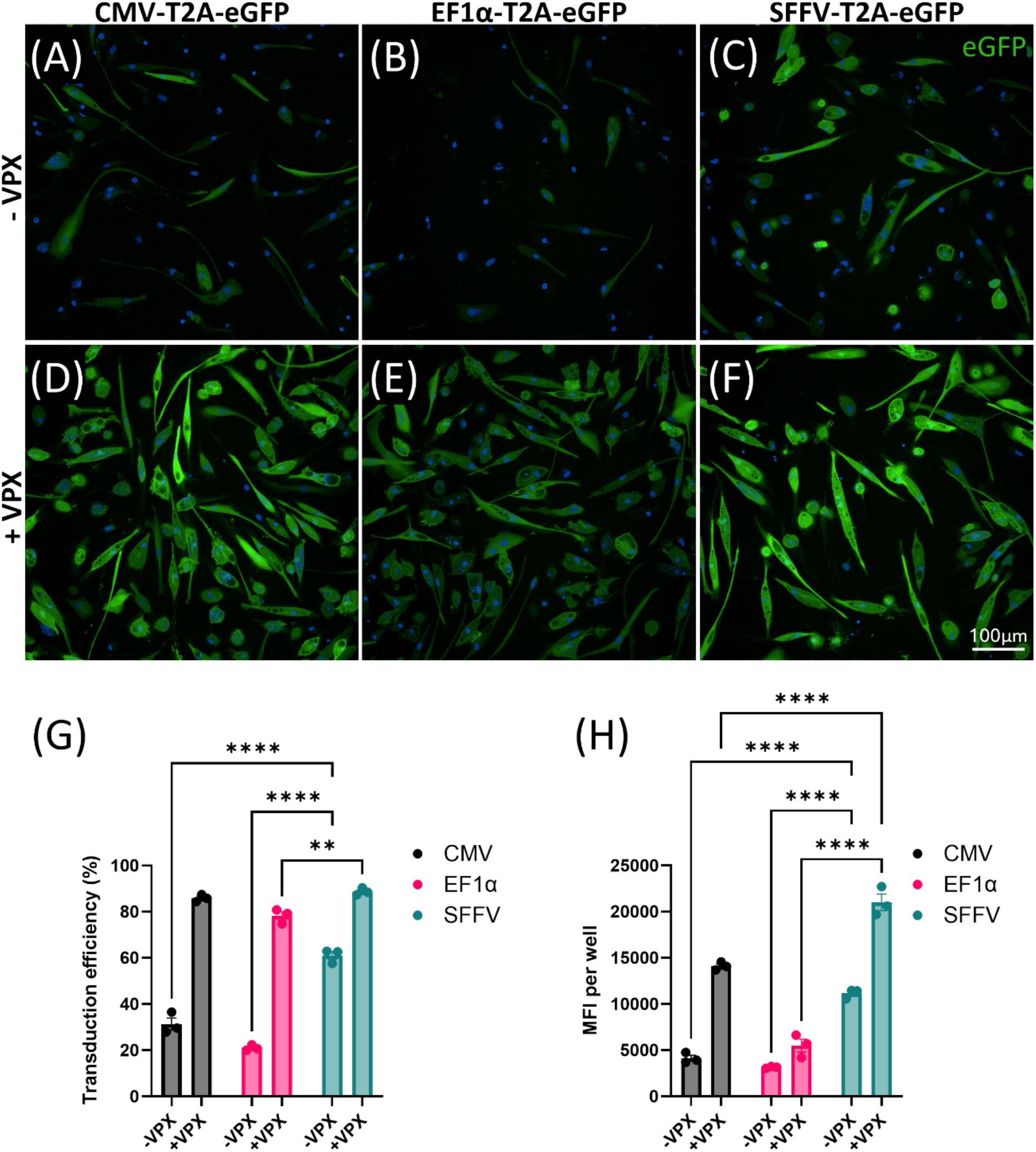
Effect of SFFV promoter in isolation in macrophages. Representative images of KOLF2.1 iPSC-derived macrophages transduced with **(A)** CMV-T2A-eGFP, **(B)**EF1α-T2A-eGFP, or **(C)** SFFV-T2A-eGFP alone and with VPX and **(D)** CMV-T2A-eGFP, **(E)** EF1α-T2A-eGFP or **(F)** SFFV-T2A-eGFP at DIV0; scale bar represents 100µm. Quantification of **(G)** transduction efficiency and **(H)** MFI at DIV10. Statistical significance was tested by two-way ANOVA (effect of promoter and VPX p=<0.0001) with Tukey’s post hoc test **p<0.005; ****p<0.0001; n=3 technical replicates per condition; error bars denote mean ±SEM.

To ensure that all components of the transduction protocol were optimised, the capability of VPX was investigated. VPX acts to downregulate SAMHD1 which normally inhibits reverse transcription of lentiviral RNA. After 24 hours, SAMHD1 expression showed a significant almost tenfold downregulation (Figure 3A, B). We therefore treated cells with VPX at the same time as applying the lentiviruses (+VPX), as previously described, or as a 24-hour pretreatment before applying the lentiviruses of interest (+24-hour VPX). There were no significant differences in transduction efficiency between these two VPX conditions. However, when combined with the SFFV promoter there was a 39% increase in MFI between +VPX and +24-hour VPX treatments. Altogether, this suggests that although SAMHD1 expression takes 24 hours to be visibly downregulated, transduction with the lentivirus of interest can occur simultaneously rather than 24 hours later for maximum transduction efficiency as the lentivirus was incubated for 48 hours total. Further transgene expression, as measured by MFI, can be attained only when using the SFFV promoter by staggering the addition of the lentivirus 24-hours after adding VPX.

For microglia, the SFFV promoter remained the most effective, providing significantly higher transduction efficiency when used alone or when combined with VPX compared to both the CMV and EF1α promoters under the same conditions. However, using the standardised plasmid backbone with cPPT and WPRE increased transduction efficiency for both CMV and EF1α promoters increasing from 6% to 41% and from 10% to 17% respectively in microglia. It was also apparent that when transduced alone the EF1α promoter performed the worst in terms of transduction efficiency and MFI (Figure 3 L, M).

To understand the impact of transducing microglia with a single lentivirus alone or in addition with VPX on microglial function we performed a pHrodo zymosan phagocytosis uptake assay. This demonstrated that VPX alone showed a significant increase in zymosan uptake suggesting a more activated microglial state (Supplemental Figure 3). However, transduction with SFFV lentivirus (plasmid F) alone or in combination with VPX showed similar levels of uptake as untreated iPSC-derived microglia. This may be an important consideration for some downstream phenotypic assays.

### The SFFV promoter increases transduction efficiency in macrophages

Next, we investigated if these improvements extend to iPSC-derived macrophages. Macrophages were transduced with plasmids F, G or H alone or with VPX delivered simultaneously with the lentivirus of interest (Figure 4). Given the results above we did not test VPX addition 24 hours prior to the addition of lentivirus. In the absence of VPX the SFFV promoter provided significantly higher transduction efficiency at 61% compared to the CMV (31%) and EF1α (21%) promoters (Figure 4G). When combined with VPX, transduction efficiency was further improved for all promoters with an increase to 89% for SFFV which outperformed both CMV (86%) and EF1α (78%). Additionally, combining these plasmids with VPX increased MFI for CMV and SFFV plasmids but this was not the case for eGFP driven by the EF1α promoter. This finding was also observed in the microglial cells. In macrophages, the SFFV promoter treatment gave the highest MFI, with and without VPX (Figure 4H).

## Discussion

With the increased appreciation of the importance of microglia in health and disease, a simple and efficient protocol to manipulate their gene expression is essential for the neuroscience field. However, both microglia and macrophages are refractory to genetic manipulation, in part due to their innate function as immune cells, and it has proven difficult to achieve high transduction efficiencies with viral vectors. Here, we demonstrate notable improvements to lentiviral vector design, specifically driving transgene expression from the SFFV promoter alongside the use of cPPT and WPRE sequences to enhance gene expression. The advantage of this approach is that high efficiency transduction can be achieved after exposure of cells to a single virus. Current methods require VPX to enhance expression, however the use of multiple viruses (including VPX) has unintended side effects such as activation, increased cell death (37) and increased rates of phagocytosis (Supplemental Figure 3). Depending on experimental requirements the complementation of the SFFV promoter with cPPT and WPRE sequences is sufficient to significantly enhance current transduction protocols for both microglia and macrophages reaching up to 73% and 61% transduction efficiency respectively with a single virus. Controlled experiments with alternate promoters (CMV or EF1α), confirmed the significant improvement of using the SFFV promoter, which can be further enhanced when combined with VPX. Therefore, should extremely high transduction efficiency and high transgene expression be required we show that using the SFFV promoter with cPPT and WPRE elements in combination with VPX treatment can provide >90% transduction and increased transgene expression. However, as shown in Supplemental Figure 3, VPX can activate iPSC-derived microglia which others have shown to hold true in monocytes (37) and so this should be considered when selecting the most appropriate transduction method.

The SFFV promoter has previously been demonstrated to be effective for *in vivo* transduction of myeloid cells (25,26,28,29). In examining the impact of different promoters in driving expression in these myeloid lineages, we show the eukaryotic promoter EF1α to be less effective than CMV in both microglia and macrophages. Although the CMV promoter is widely used across many applications previous work has suggested that it is not well suited to iPSC-based experiments where the promoter is often silenced (16–18). Our findings have also highlighted the importance of the plasmid backbone. From the work above, the effect of both the WPRE and cPPT elements are apparent in improving transduction efficiency alongside streamlined plasmid design. WPRE is thought to enhance target gene expression through post-transcriptional regulation whilst cPPT has been reported to improve gene transcription and vector integration (32–36).

## Conclusion

With the increased interest and requirement to modify microglial gene expression, and that of related cell types such as macrophages, our findings provide an improved tractable approach enabling effective lentiviral transduction that can be readily applied to a wide range of experimental paradigms. This includes: simple gene expression manipulation for disease-modelling, including *in vitro* models of neurodegenerative diseases; expression of reporter proteins, such as GFP, that can be used for easy tracking of these cell types (38) in coculture, organoids or in xenotransplantation models and for screening purposes; as well as for the introduction of optogenetic or chemogenetic tools to manipulate and examine microglial activity. Overall, this work will enable easier investigation of microglial and macrophage biology across the neuroimmune landscape.

## Declarations

### Availability of data and materials

The datasets used and/or analysed during the current study are available from the corresponding author on reasonable request. Plasmids will be shared upon request.

### Author Contributions

SG, PRT and NCR designed experiments, SG and MAC performed experiments, SET provided macrophage precursors, SG collected and analysed the data and produced the figures, NCR conceived the project, SG and NCR wrote the manuscript, NCR supervised the work. All authors read and approved the final text.

## Acknowledgements

We thank Prof Vincent Dion for sharing the CMV-eGFP /EF-1α-mCherry lentivirus (Plasmid A). Supplemental Figure 3 was created in BioRender. Goberdhans, S. (2026) https://BioRender.com/gk4wq7a

