## Supplemental Figures for "Highly Efficient Lentiviral Transduction of Human iPSC-Derived Microglia and Macrophages"

### Supplemental Figures

(A)

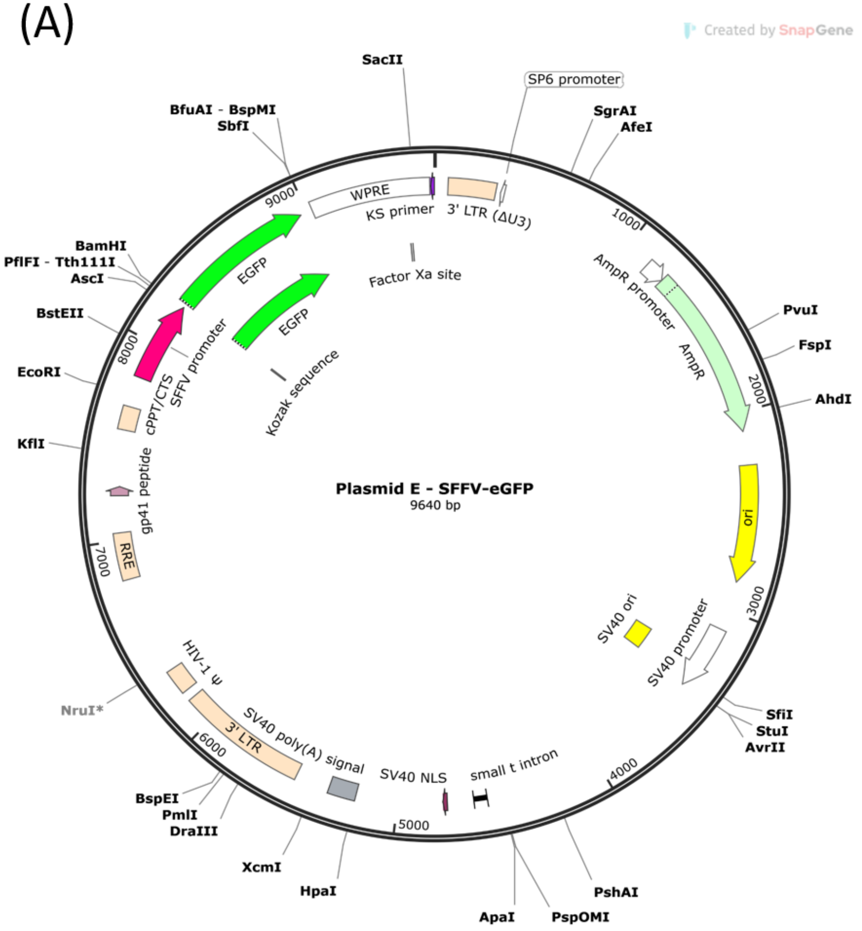


(B)

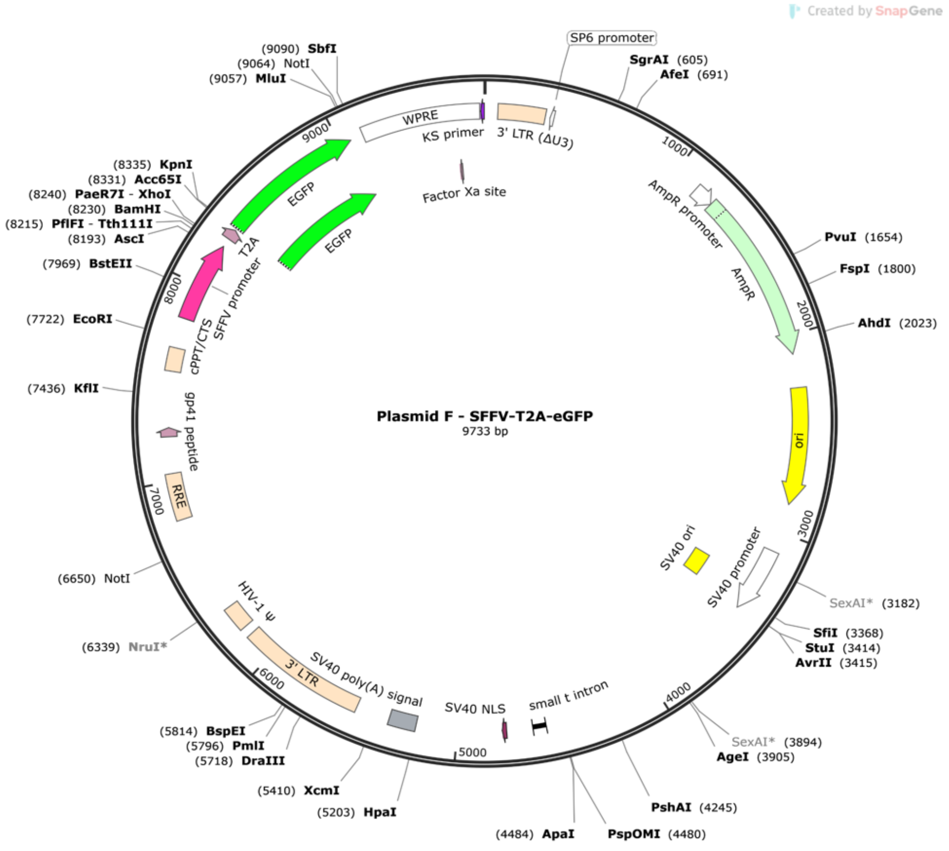

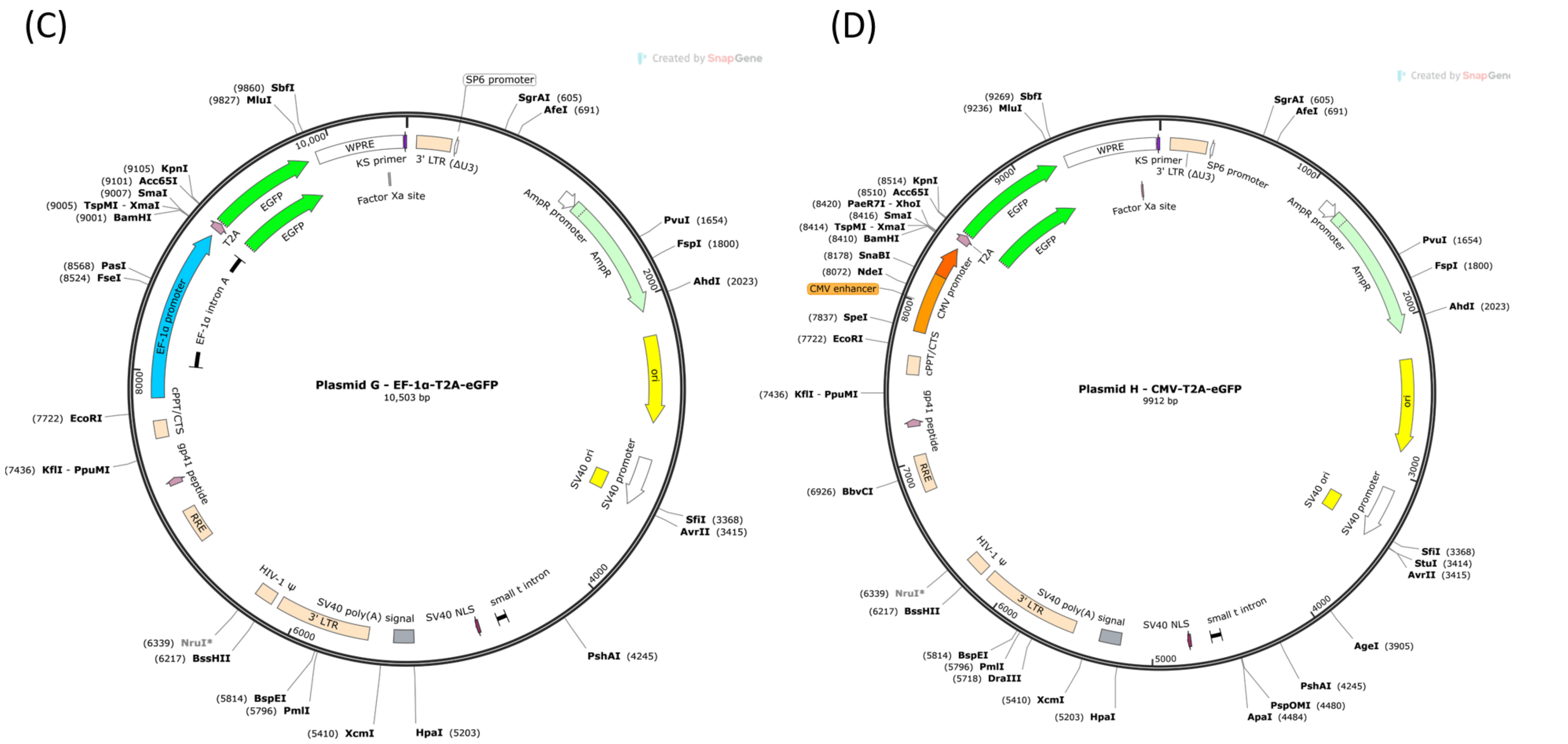


***Supplemental Figure 1: Plasmid maps***

***(A)*** *Plasmid map for Plasmid E, SFFV-eGFP used in Figure 2.* ***(B-D)*** *All four plasmids have the same backbone with only the promoter region modified* ***(B)*** *Plasmid F (SFFV-T2A-eGFP)* ***(C)*** *Plasmid G (EF1α-T2A-eGFP) and* ***(D)*** *Plasmid H (CMV-T2A-eGFP).*

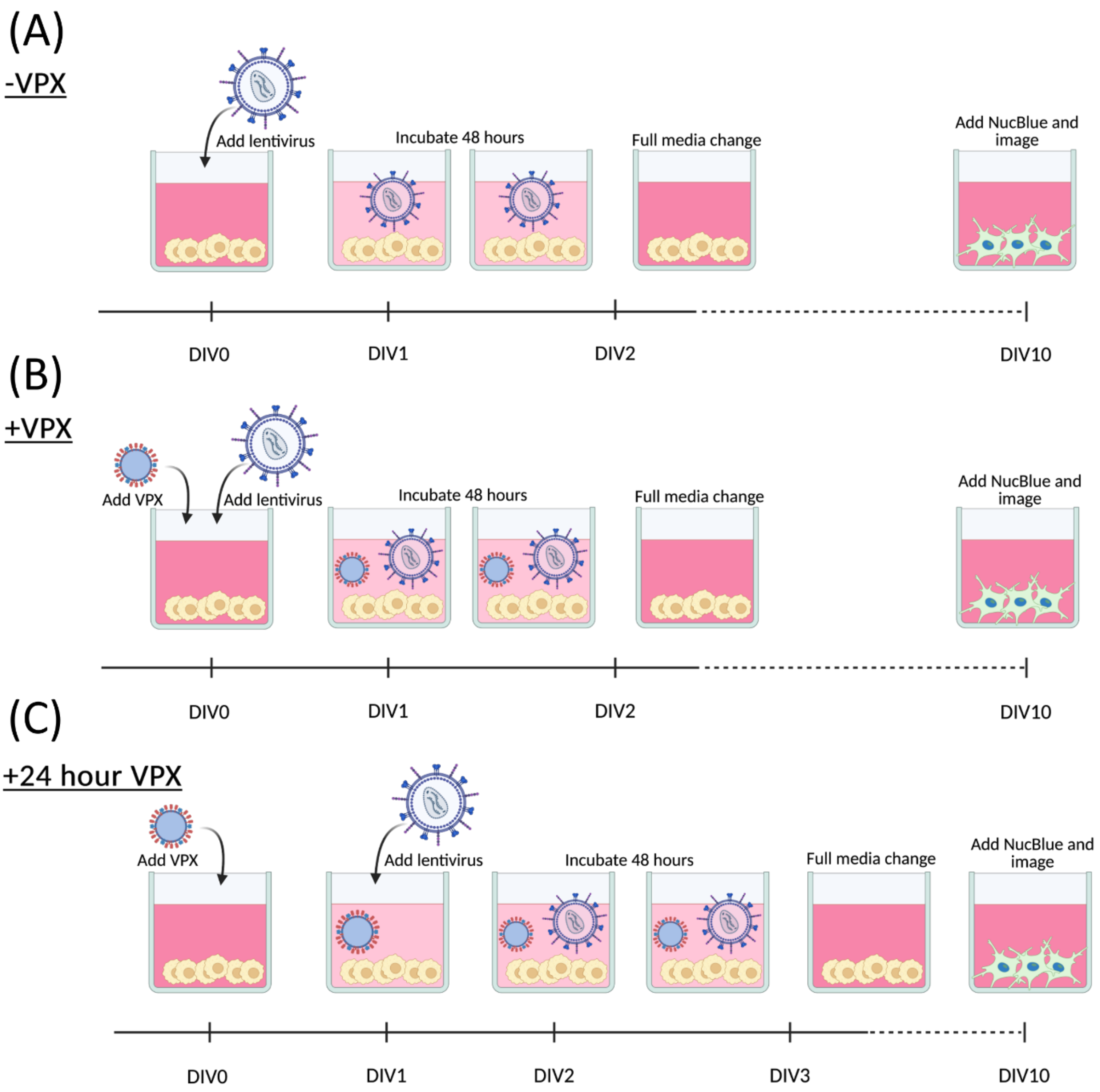


***Supplemental Figure 2: Lentiviral transduction timeline***

***(A)*** *Transduction of cells without VPX, lentivirus added at DIV0 and incubated for 48 hours prior to full media change*

***(B)*** *Transduction of cells with VPX, both lentivirus and VPX added at DIV0 and incubated for 48 hours prior to full media change*

***(C)*** *Transduction of cells with 24-hour pretreatment of VPX, VPX added at DIV0, 24 hours later lentivirus was added and incubated for 48 hours prior to full media change.*

*All cells matured until DIV10, nuclei stained with NucBlue and imaged using the Opera Phenix.*

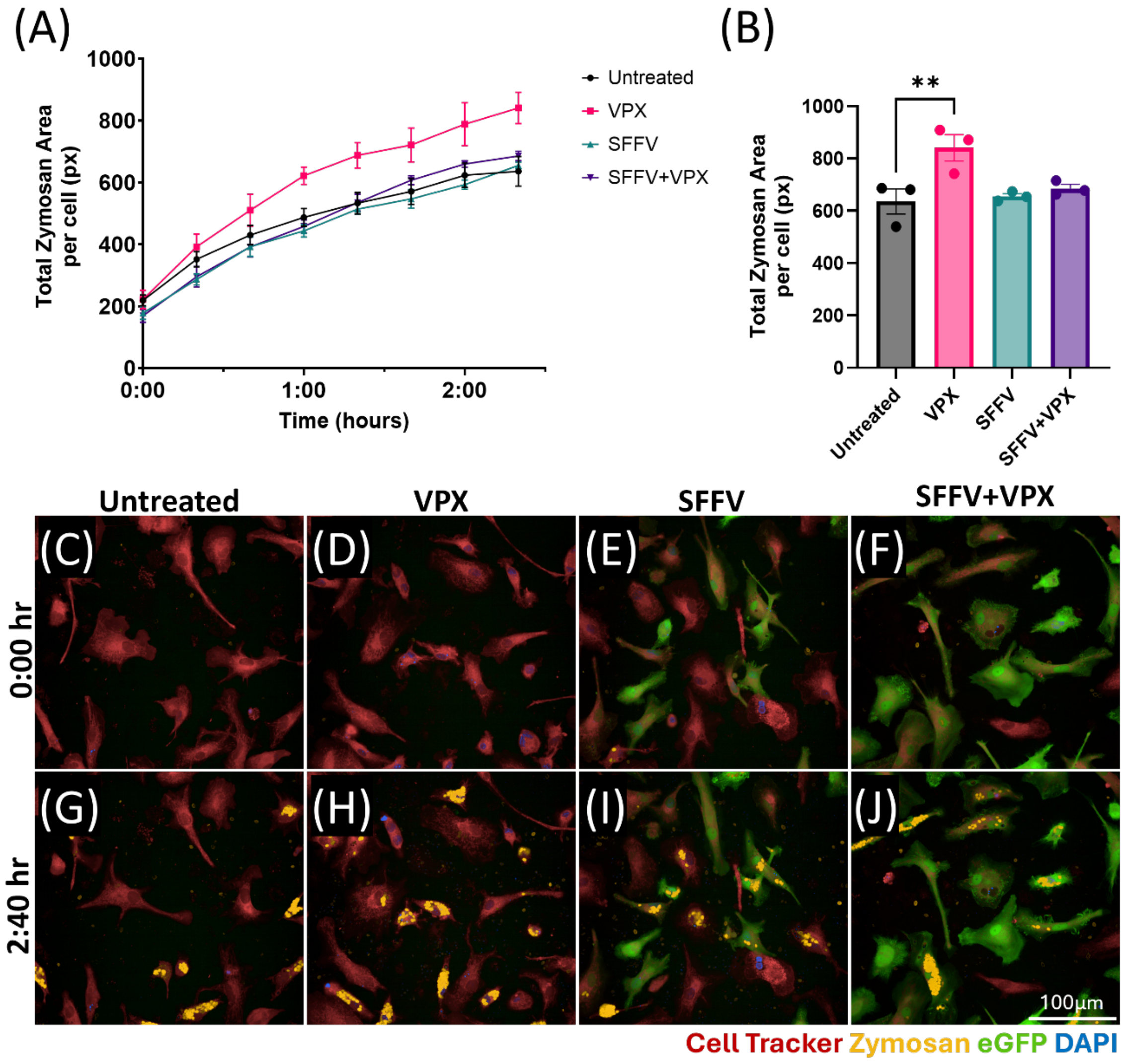


***Supplemental Figure 3: Effect of lentivirus transduction on microglial function***

***(A)*** *Zymosan uptake normalised to total DAPI positive cells in DIV10 microglia alone (untreated) or microglia transduced with VPX only, SFFV only, or SFFV and VPX, imaged at intervals of 20 minutes.* ***(B)*** *Quantification of the final timepoint with statistical significance tested by one-way ANOVA (p<0.05) with Dunnett's post hoc test with the untreated condition as the control **p <0.01 Representative images of microglial uptake of pHrodo zymosan particles from each condition at* ***(C-F)*** *0 hours and* ***(G-J)*** *after 2 hours 40 minutes; scale bar represents 100μm*
